# Exact distributions of threshold crossing times of proteins under post-transcriptional regulation by small RNAs

**DOI:** 10.1101/2024.08.05.606600

**Authors:** Syed Yunus Ali, Ashok Prasad, Dibyendu Das

## Abstract

The timings of several cellular events like cell lysis, cell division, or pore formation in endosomes are regulated by the time taken for the relevant proteins to cross a threshold in number or concentration. Since protein synthesis is stochastic, the threshold crossing time is a first passage problem. The exact distributions of these first passage processes have been obtained recently for unregulated and auto-regulated genes. Many proteins are however regulated by post-transcriptional regulation, controlled by small non-coding RNAs (sRNAs). Certain mathematical models of gene expression *with* post-transcriptional sRNA regulation have been recently exactly mapped to models *without* sRNA regulation. Utilizing this mapping and the exact distributions, we calculate exact results on fluctuations (full distribution, all cumulants, and characteristic times) of protein threshold crossing times in the presence of sRNA regulation. We derive two interesting predictions from these exact results. We show that the size of the fluctuation of the threshold crossing times have a non-monotonic U-shaped behavior as a function of the rates of binding and unbinding of the sRNA-mRNA complex. Thus there are optimal parameters that minimize noise. Furthermore, the fluctuations in models with sRNA regulation may be higher or lower compared to the model without regulation, depending on the mean protein burst size.

## I. INTRODUCTION

Many molecular processes in cells have been shown to behave as if they are controlled by an internal clock. For example, in a linear sequence of enzymatic modifications, enzymes are often found to be synthesized in temporal order. Thus, enzymes that work later in the pathway are produced after the enzyme catalyzing the previous step^1^. Another example is cell lysis after lambda-phage infection, which appears to be the result of an evolutionary process to optimize the survival of the virus: too long and the virus may die with the bacteria, too soon and not enough virions are produced^2,3^. An attractive and natural hypothesis for the mechanism of determining the timing for such processes is that it is determined by the time taken for the number or concentration of one or more proteins reaching a threshold^1,4–6^.

This timing mechanism is intrinsically stochastic because gene expression itself is inherently stochastic. Synthesis of mRNA by transcription of DNA and synthesis of proteins through translation are both intrinsically stochastic processes^7–15^. These stochastic events produce different quantities of mRNA and proteins in different cells, even if they are genetically identical and exposed to the same external conditions^16,17^. Given this stochasticity, the time to reach a protein threshold is a *first passage time* problem^18^. This first passage time has been shown to be important in several biological contexts such as cell lysis on infection of E.coli by bacteriophage-*λ* via threshold crossing of the protein Holin^2,19^, pore formation in endosomes enclosing Streptococcus Pneumonia and threshold crossing of toxin Pneumolysin^20^, cell division in bacteria like E.coli and Caulobacter^21–23^ and cell cycle time regulation with threshold crossing of relevant proteins^24–26^.

The corresponding theoretical first passage problem has been solved for its moments^27^ in the bursty protein production limit, and recently for the full distribution using stochastic formalism in backward time^5,6^. These previous papers however studied cases where the gene of interest is either not regulated or positively or negatively auto-regulated. But post-transcriptional regulations are ubiquitous in both prokaryotes and eukaryotes, for which no such analytical results exist.

In this paper we present new results for proteins that are regulated through post-transcriptional regulation by small non-coding RNAs. Considerable research underscores the critical role of noncoding RNAs in regulating gene expression^28–33^. A significant focus has been on small ncRNAs or simply small RNAs (sRNAs). The sRNAs bind to mRNAs and control protein production by altering mRNA stability or regulating translational efficiency^34–36^. Many sRNAs are known to repress gene expression, thus regulating it negatively^37–41^. Sometimes, sRNAs can also activate or switch between activation and repression depending on cellular signals; thus, positive regulation is also possible^39,42,43^. Post-transcriptional regulation is pivotal in various cellular processes, from stress responses^44^ to the expression of virulence genes^45–48^. Interestingly, it has been shown that the timing of the iron stress response in a cyanobacterial organism is controlled by small RNAs^49^. Small RNAs also appear to play an important role in controlling the timing of developmental events^50^. Thus the stochastic properties of sRNA controlled timing events should be very useful for cell biology.

Experimental techniques such as single-cell RNA sequencing (scRNA-seq) and single-molecule fluorescence in situ hybridization (smFISH) to quantify mRNA, and fluorescent proteomic imaging, mass cytometry and mass spectrometry to quantify protein levels in individual cells^51–55^ have been used to demonstrate the stochastic nature of mRNA and protein production. A major source of noise in mRNA levels is transcriptional bursting, arising due to the promoter switching between transcriptionally active and inactive states^56,57^. On the theoretical side, the full joint distribution of mRNA and protein with such telegraphic switching of the promoter has been recently exactly solved^58^. It is known that often in bacteria and yeast, protein production arises in short intense bursts, which are geometrically distributed^13,59^. These bursts may arise from multiple ribosomes translating the same mRNA, rapidly decay of short-lived mRNA leaving behind long-lived proteins^14,15,60,61^, or due to switch between translationally active and inactive states^62^. In this paper we would focus on this limit, where protein production occurs effectively in bursts.

A mathematical formalism for post-transcriptional regulation by small RNA (sRNA) has been developed^39,63–66^. Some recent studies have theoretically studied the statistics of first passage times (FPT) in certain models of sRNA-mediated regulation^67–69^, by finding an exact mapping of the problems with regulation to problems without regulation. Their analysis, nevertheless, was approximate. In previous works we reported exact analytical results on the FPT distribution for the unregulated problem^5,6^. Based on these results, and the mapping mentioned above, here we derive exact results for the full FPT distribution and measures of fluctuation associated with it, for proteins regulated by sRNAs. We carry out Kinetic Monte Carlo simulations and show an accurate match with the analytical expressions, unlike in the previous literature. Moreover, we show that from the variation of metrics of fluctuation (coefficient of variation *CV*, Skewness, and relative characteristic times) on the parameters of the unregulated model, we can derive the exact dependence of such metrics on parameters of the models with sRNA regulation. We find a generic U-shape in the metrics of fluctuation as a function of parameters related to sRNA regulation. The relative fluctuations of the regulated and unregulated models can also now be compared exactly.

The paper is organized as follows. In Sec. II, we introduce the two-stage model of gene expression without regulation, discuss the bursty protein production limit, and the exact distribution of first passage times in that limit. Next, we present the different models of gene expression with sRNA regulation and their exact mapping to the unregulated model. In Sec. III we present various exact results on the fluctuations of FPTs in the sRNA regulated models, and compare those with KMC simulations. We present the concluding remarks in Sec. IV.

## II. MODEL SUMMARY

We describe the models of gene expression that we study, in Fig. 1. For these models, we would study the statistical properties of the first passage times, for the protein number to reach a threshold *X* for the first time.

**FIG. 1:**
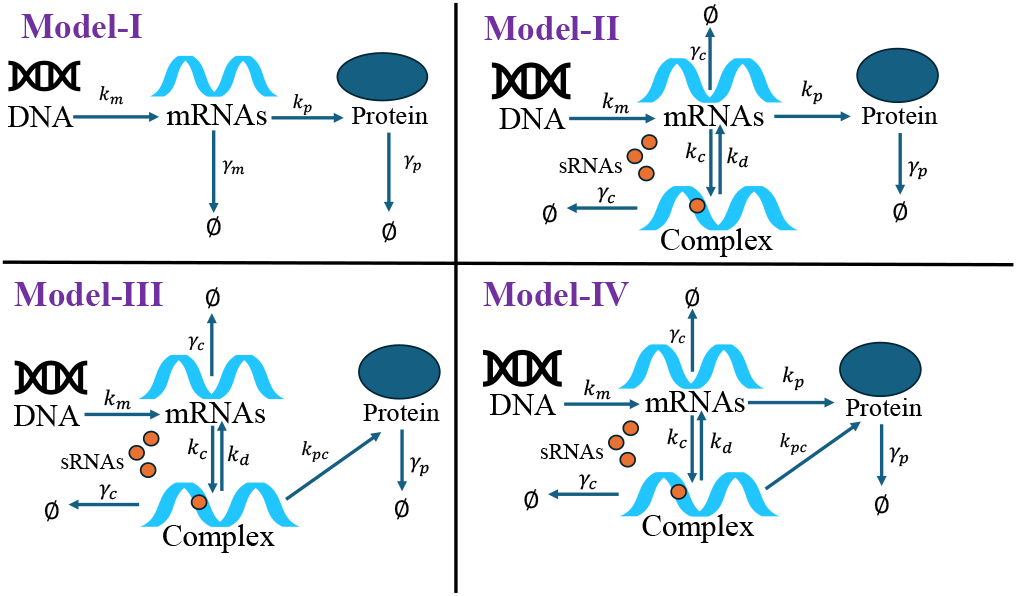
Schematic representation of the models of gene expression. For all the models, the transcription rate is *k*_*m*_, the translation rate (from mRNAs) is *k*_*p*_, and the degradation rates of mRNAs and proteins are *γ*_*m*_ and *γ*_*p*_, respectively. Model-I: Basic two-stage model without transcriptional regulation. Post-transcriptional regulation modifies Model-I in the remaining three models. Model-II: mRNA and sRNA associate to form a complex with rate *k*_*c*_, and the complex dissociates back at rate *k*_*d*_. Model-III: mRNAs do not translate directly. Translation of protein happens from mRNA-sRNA complex at rate 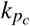. Model-IV: Translational happens from both mRNA and mRNA-sRNA complex. The degradation rate of mRNA-sRNA complexes is *γ*_*c*_ for Models II, III, and IV.

### A. Model-I

This is the basic two-state model without regulation, where DNA is transcribed to mRNAs at rate of *k*_*m*_ and mRNAs translated to proteins at rate *k*_*p*_. The mRNAs and proteins degrade at rates *γ*_*m*_ and *γ*_*p*_, respectively. A schematic representation is shown in Fig. 1. At any time t, let *m* and *n* be the number of mRNAs and proteins, respectively, with probability *P*(*m, n, t*). This probability evolves following the Master equation^13^:

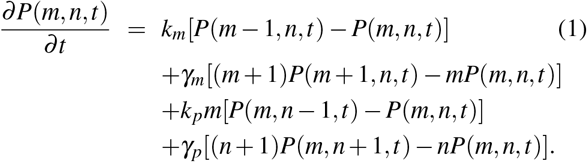

The equation above is non-dimensionalized by introducing parameters 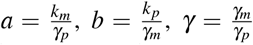 and *τ* = *γ*_*p*_*t*. Here, *a* represents the mean number of mRNA bursts per cell cycle, *b* represents the mean number of protein molecules produced from an mRNA per burst^59^, and *γ* is the number of mRNA molecules that degrade in the time that one protein molecule degrades. By defining the generating function, *G*(*x, z, t*) = ∑_*m,n*_ *x*^*m*^*z*^*n*^*P*(*m, n, t*) with *x* = *u* + 1 and *z* = *w* + 1, Eq. 1 leads to the partial differential equation^13^:

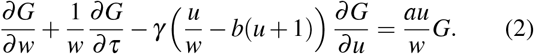

Under the assumption that the protein lifetime is much longer than the mRNA lifetime (*γ*≫1), which is often found in bacteria, yeast and mammalian cells^13,59,70^, following the method of Lagrange characteristics, it may be shown that the generating function obeys^13^:

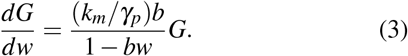

Moreover in this limit (*γ*≫ 1), proteins have a *bursty* production – effectively a burst of size *r* from a single mRNA translation follows a geometric distribution 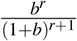^13^. The effective Master equation for the probability *P*(*n, τ*) of *n* proteins is

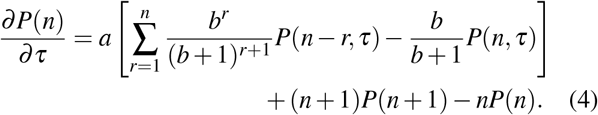

Just like the forward Master Eq. 4 helps to study the stochastic protein number (*n*) distribution at any chosen time *t*, the statistics of the first passage times may be studied through the following backward Master equation (BME) in the *bursty* protein production limit^5^.

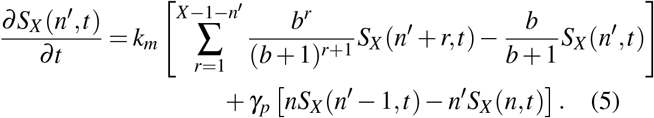

Here, the survival probability *S*_*X*_ (*n*^′^, *t*) is the probability that the protein number, initially starting from *n*^′^ at *t* = 0, stays below the threshold *X* up to time *t*. There is an initial condition that *S*(*n*^′^ *< X*, 0) = 1, and an absorbing boundary condition *S*(*n*^′^ ≥ *X*) = 0. The probability distribution of first passage times *f*_*X*_ (*n*^′^, *t*) is obtained from the survival probability as follows:

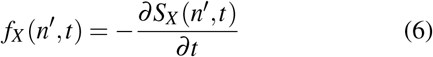

One may define Laplace transforms 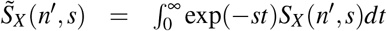 and 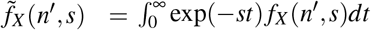 for the survival probability and first passage time distribution respectively.

In the recent study by Rijal et al. (2020)^5^, the BME (Eq. 5) was exactly solved in Laplace space. In particular, the Laplace transform of the first passage distribution starting with zero number of proteins (*n*^′^ = 0) is given by:

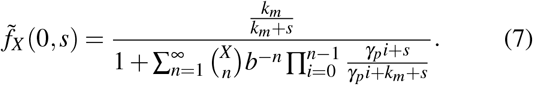

#### 1. Utility of Eq. 7

The exact expression in Eq. 7 is useful in many ways, which we discuss below– we will extensively use these results in the paper.

I. Firstly, one may obtain the distribution in the time domain by numerically inverting the Laplace transform in Eq. 7 using Mathematica^71^. This exact distribution *f*_*X*_ (0, *t*) may then be compared to distributions obtained through the Kinetic Monte Carlo (KMC) simulations. We do this for different models in this paper.
II. Secondly, Eq. 7 may be used to obtain the moments of the first passage time of *m*-th order:

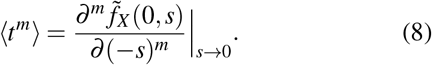 The explicit expressions of the second and third moments are available in^5,27^, but Eq. 8 can also be evaluated using Mathematica. These are useful for obtaining the coefficient of variation *CV*^2^ and Skewness:

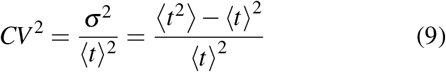

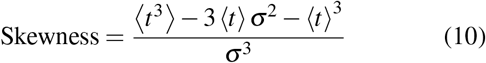
III. The distribution of first passage times *f*_*X*_ (0, *t*) has an exponential tail ∼ exp(− *t/τ*_*c*_), where *τ*_*c*_ is called the characteristic time^6^. The pole in the denominator of Eq. 7, whose real part − *α*_*c*_ is closest to the origin in the complex *s*-plane, gives the characteristic time:

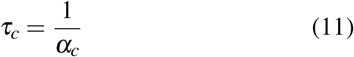

One may determine the roots of the denominator of Eq. 7, identify *α*_*c*_, and thus determine *τ*_*c*_.

#### 2. Details of KMC simulations

To compare with our analytical predictions, we perform Kinetic Monte Carlo (KMC) simulations of the Models-II, III & IV following the standard Gillespie algorithm^72^. Given a state of the system in any instant of time, one may identify a list of possible events which would change it. If these events have a set of rates {*r*_*i*_ }, any one of them may be chosen according to the probabilities *r*_*i*_*/R* where *R* = ∑_*i*_ *r*_*i*_. After the event is accepted and the system is updated, the time *t* is incremented to *t* + δ*t*, where δ*t* is distributed as *P*(δ*t*) = (∑_*i*_ *r*_*i*_) exp(−δ*t* ∑_*i*_ *r*_*i*_), as expected in a Poisson process. The simulation is stopped when *n*(*t*_*FPT*_) = *X* for the first time and *t*_*FPT*_ gives us the desired time of first passage. Subsequently such randomly sampled times over repeated histories help us calculate various statistical measures of FPT fluctuations. The number of sample FPT used in KMC simulations for every data point in the Figures below is 10^7^.

#### 3. Characteristic times from extreme statistics

Although characteristic times can be theoretically calculated using Eq 11, it is also possible to be estimated from numerically sampled FPTs. This is done by using the statistics of extreme times in subsets of the data. It is well known that extremes (maxima of data sets) for exponential-tailed distributions follow a Gumbel distribution and its variance *σ*^2^ is related to the characteristic time *τ*_*c*_ of the original distribution as^73^

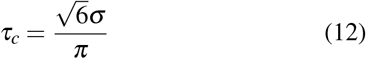

The above formula may be used to obtain *τ*_*c*_ from KMC simulation data, as done in some past literature^74^. If there are *N*× *M* sampled times, the data may be split into *M* sets, each with *N* sample times. Each of the *M* sets provides a maximum *t*_*max*_ such that there would be *M* such values. These *t*_*max*_ values may then be used to obtain *σ*, which leads to *τ*_*c*_ through Eq. 12. In typical KMC simulations, we used *M* = 10, *N* = 10^5^ and 200 such histories to obtain the averaged *τ*_*c*_.

In all three models II, III and IV, that we discussed below we assume that the sRNA regulators are abundant enough that their binding or unbinding to a single mRNA target does not significantly alter the free sRNA concentration.

### B. Model-II: Negative Post-transcriptional Regulation

One class of prokaryotic sRNAs regulates protein production after transcription by associating with mRNAs and forming a sRNA-mRNA complex. The complex does not actively produce proteins, and hence, the sRNA binding suppresses protein production. This is refer to as negative post-transcriptional regulation^39,68,69,75,76^. A Schematic diagram of this scenario is given in Fig. 1. All rates are like Model-I; additionally, mRNA forms mRNA-sRNA complex at rate *k*_*c*_, and the complex dissociates at rate *k*_*d*_. Similar to the quantity *γ* defined for Model-I, in this model, we may define *γ*′ = *γ*_*c*_*/γ*_*p*_, the ratio of degradation of sRNA-mRNA complex to protein. The configuration of the system is specified by the number of mRNAs *m*, number of proteins *n*, and number of mRNA-sRNA complex *m*_*c*_, which has a probability *P*(*m, m*_*c*_, *n, t*). The Master equation evolving this probability is shown in the appendix. A.

It is interesting that in a certain limit this model is exactly mapped to Model-I. Following^69^ and assuming *γ*_*m*_ = *γ*_*c*_ and *γ*′(= *γ*) ≫1 (*bursty* protein production limit) it may be shown (appendix. A) that the generating function of Model-II becomes identical to that of Model-I given by Eq. 3, with *b* being replaced by

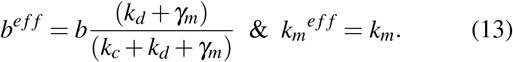

This is a very powerful mapping which we use in this paper to predict the properties of the Model-II (with post-transcriptional regulation) by varying parameters *k*_*c*_ and *k*_*d*_, in terms of exactly known properties of Model-I (without regulation). In particular this will help in studying the fluctuations of first passage times in Model-II.

### C. Model-III: Positive Post-transcriptional Regulation

As discussed in^39^ certain mRNA structures may be such that ribosomes cannot ordinarily bind and produce proteins. But sRNA binding to the mRNA exposes ribosome binding sites and facilitates translation. This is called positive post-transcriptional regulation^39,67,77^. This scenario is effectively described by Model-III, which has no direct translation from mRNAs but translation from the mRNA-sRNA complexes at rate 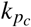 (Fig. 1). The rest of the rates are as in Model-II. The Master equation evolving with probability *P*(*m, m*_*c*_, *n, t*) is shown in the appendix B^67^.

This model may also be exactly mapped to Model-I, under the assumptions *γ*_*m*_ = *γ*_*c*_ and *γ*′(= *γ*) ≫ 1 (*bursty* protein production limit). That means that the equation of the generating function *G* of Model-III in this limit is given by Eq. 3, with *b* and *k*_*m*_ being replaced by following effective values:

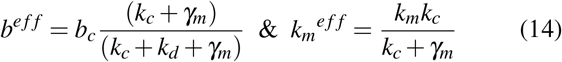

Here *b*_*c*_ = *k*_*pc*_*/γ*_*p*_. Hence, one may predict the properties of Model-III on the variation of parameters *k*_*c*_ and *k*_*d*_ based on properties known for Model-I. Specifically, we will utilize this mapping to study the fluctuations of protein threshold crossing times in Model-III.

### D. del-IV: Combining Models-II and III

Model-IV represents a generalized process that combines both Model-II and Model-III. In this model, protein translation occurs from both mRNAs and mRNA-sRNA complexes, with rates *k*_*p*_ and *k* 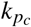 respectively. The schematic diagram of this model is illustrated in Fig. 1. The rates in Model-IV are consistent with those in Models I, II, and III. The Master equation for this model can be found in the appendix C, following the approach outlined in previous studies^63,69^. As in Model-II and Model-III, we would proceed with the assumptions *γ*_*m*_ = *γ*_*c*_ and *γ*′(= *γ*) *>>* 1 (*bursty* protein production limit). But that merely does not make the model map to Model-I as in the other two cases. One has to make a further assumption that the burst sizes of proteins produced from both mRNAs and mRNA-sRNA complexes are the same (*b* = *b*_*c*_). Essentially, the distinction between mRNAs and mRNA-sRNA complexes vanish under these assumptions, leading to the conclusion that the generating function of the model is given by Eq. 2 with

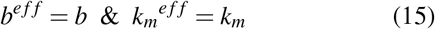

## III. RESULTS

### A. Exact distribution of threshold crossing times in models with sRNA regulations

As discussed in Section II A 1 (point number (I)), the full distribution of FPT may be derived using Eq. 7. Now, for a model with sRNA regulation, it may seem difficult *a priori* to obtain the FPT distribution. But for Model-II, using the exact mapping of effective parameters through Eq. 12 to Model-I, we actually know its exact FPT distribution. Thus for any given set of parameters of Model-II, once we calculate *b*^*e f f*^ and *k*_*m*_^*e f f*^, we put that in place of *b* and *k*_*m*_ and use all the exact results of Model-I. Similarly, in cases of Model-III and Model-IV, Eqs. 13 and 14 may be used, respectively.

To demonstrate the significance of this, we perform KMC simulations on Model-II, III, and IV for various parameters as indicated in Fig 2, but keeping their *b*^*e f f*^ and *k*_*m*_^*e f f*^ same as *b* and *k*_*m*_ of Model-I. We see that KMC FPT distributions for the models with sRNA regulation all collapse onto the exact distribution of Model-I (shown in solid line).

**FIG. 2:**
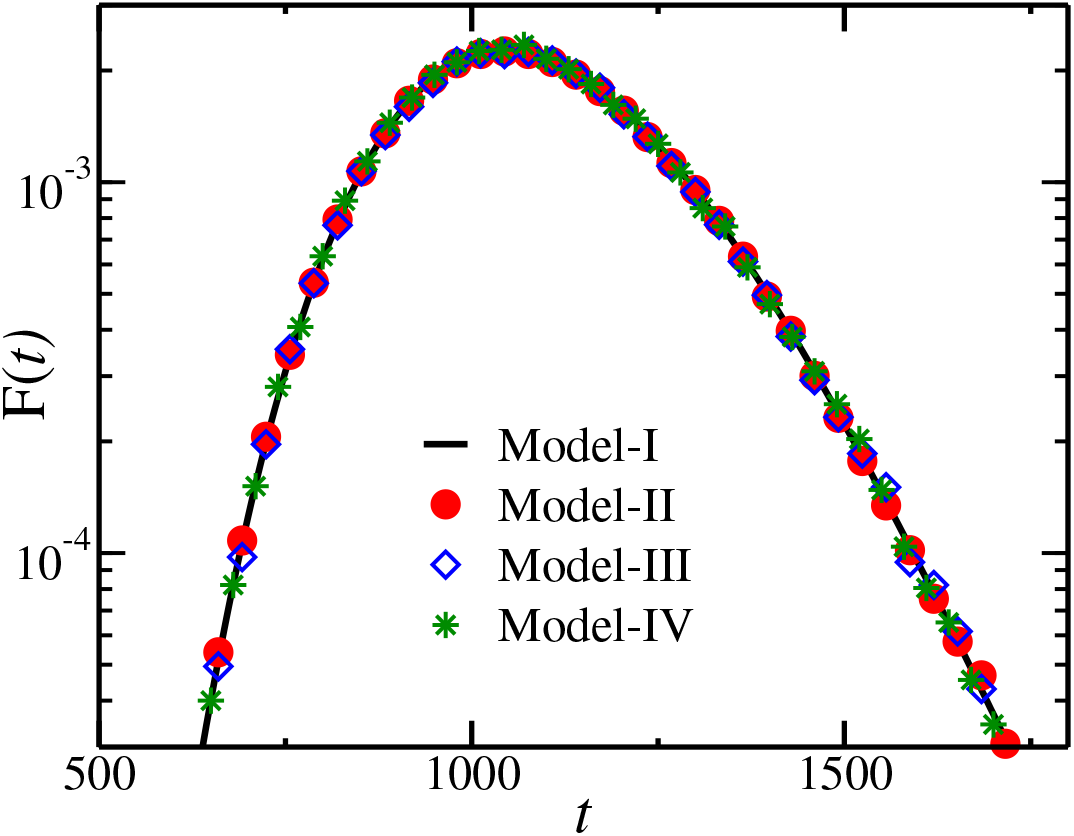
The graph shows the First-Passage Time (FPT) distributions for three different models at *X* = 400. Points are obtained from Kinetic Monte Carlo (KMC) simulation studies, and the solid line represents the exact FPT distribution. Parameters sets: Model-I: *b* = 2, *k*_*m*_ = 0.3, Model-II: *k*_*m*_ = 0.3, *k*_*p*_ = 12, *γ*_*m*_ = 6, *γ*_*c*_ = 6, *k*_*c*_ = 0.001, *k*_*d*_ = 20.0, Model-III: *k*_*m*_ = 0.39, 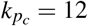, *γ*_*m*_ = 6, *γ*_*c*_ = 6, *k*_*c*_ = 20.0, *k*_*d*_ = 0.001, and Model-IV: *k*_*m*_ = 0.3, *k*_*p*_ = 12, 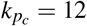, *γ*_*m*_ = 6, *γ*_*c*_ = 6, *k*_*c*_ = 10.0, *k*_*d*_ = 1. All rate parameters are in units of min^−1^.

### B. Model-I: Dependence of metrics of FPT fluctuations on burst size *b* and transcription rate *k*_*m*_

In the remaining part of the paper, we will focus on the metrics of relative fluctuations of FPT, namely *CV*^2^, skewness, and *τ*_*c*_*/* ⟨*t*⟩all of which are defined in Section II A 1 (points (II) & (III)). We would be particularly interested in their dependence on two parameters, *k*_*c*_ and *k*_*d*_ characterizing the association and dissociation of the mRNA-sRNA complex in post-transcriptionally regulated Models-II & III. But since these parameters combine to give effective *b*^*e f f*^ and *k*_*m*_^*e f f*^ through Eqs 12 and 13, we would first study these relative fluctuations for Model-I as a function of *b* and *k*_*m*_. Understanding the behaviour of the latter in the unregulated Model-I will help us understand the corresponding behaviour in regulated Models-II & III, as we would see in the subsequent sections.

It is well known that *CV*^2^ in the bursty limit of Model-I has a non-monotonic form with variation of the threshold *X* (or mean FPT)^5,6,27^. What we observe here is that as a function of burst size *b* too, *CV*^2^ decreases and then increases (see Fig 3). With increasing *X* the minima gets deeper. Moreover similar non-montonicity is present in the third order fluctuations (Skewness), and the relative characteristic time *τ*_*c*_*/* ⟨*t*⟩ (see the left panel of Fig 3). On the other hand, with transcription rate *k*_*m*_, all the three metrics of fluctuation show a monotonic decrease (see the right panel of Fig 3). We proceed below to see how these non-monotonicities reflect in Model-II and III.

**FIG. 3:**
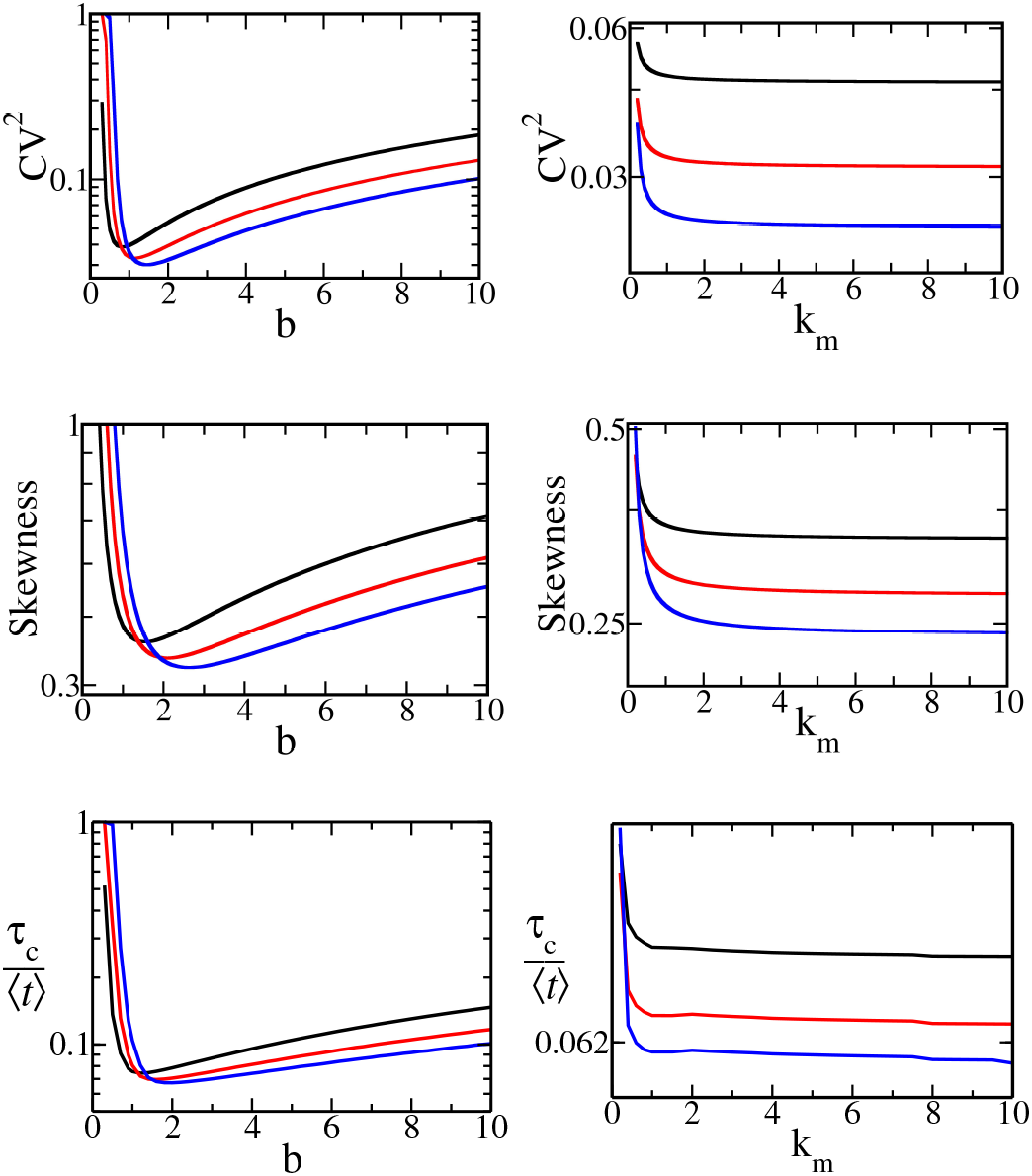
Variation of *CV*^2^, Skewness and relative characteristic times *τ*_*c*_*/* ⟨*t*⟩ with *b* (left panel, *k*_*m*_ = 0.3 min^−1^) and *k*_*m*_ (right panel, *b* = 2). Each sub-figure shows three curves for different values of protein threshold (*X*): black (*X* = 100), red (*X* = 150) and blue (*X* = 200). The protein degradation rate is fixed at *γ*_*p*_ = 0.001 min^−1^.

### C. Model-II: Variation of relative fluctuations with switching parameters *k*_*c*_ and *k*_*d*_

In the case of negative post-transcriptional regulation, we would like to study how the statistics of protein threshold crossing times depend on the model parameters *k*_*c*_ and *k*_*d*_ crucially associated with the mRNA-sRNA complex. First, we note that in Eq. 13 (for fixed *k*_*m*_), the parameter *b*^*e f f*^ monotonically decreases with *k*_*c*_, while *k*_*m*_^*e f f*^ stays unchanged. Then from the sub-figures in the left panel of Fig 3, an exact prediction immediately follows — in Model-II, the metrics of relative fluctuation will all have a non-monotonic behaviour with variation in *k*_*c*_ (as that changes *b*^*e f f*^). Armed with the theoretical curves from Model-I in the bursty limit, we compare those with the corresponding *CV*^2^, Skewness and *τ*_*c*_*/* ⟨*t*⟩ obtained using KMC simulations of the full Model-II – the match is perfect in the left panel of Fig 4. Note that theoretical *τ*_*c*_ is calculated from the pole of the Laplace transform (see Eq 11), and *τ*_*c*_ is calculated from the KMC data using the method of extremes (Eq 12).

**FIG. 4:**
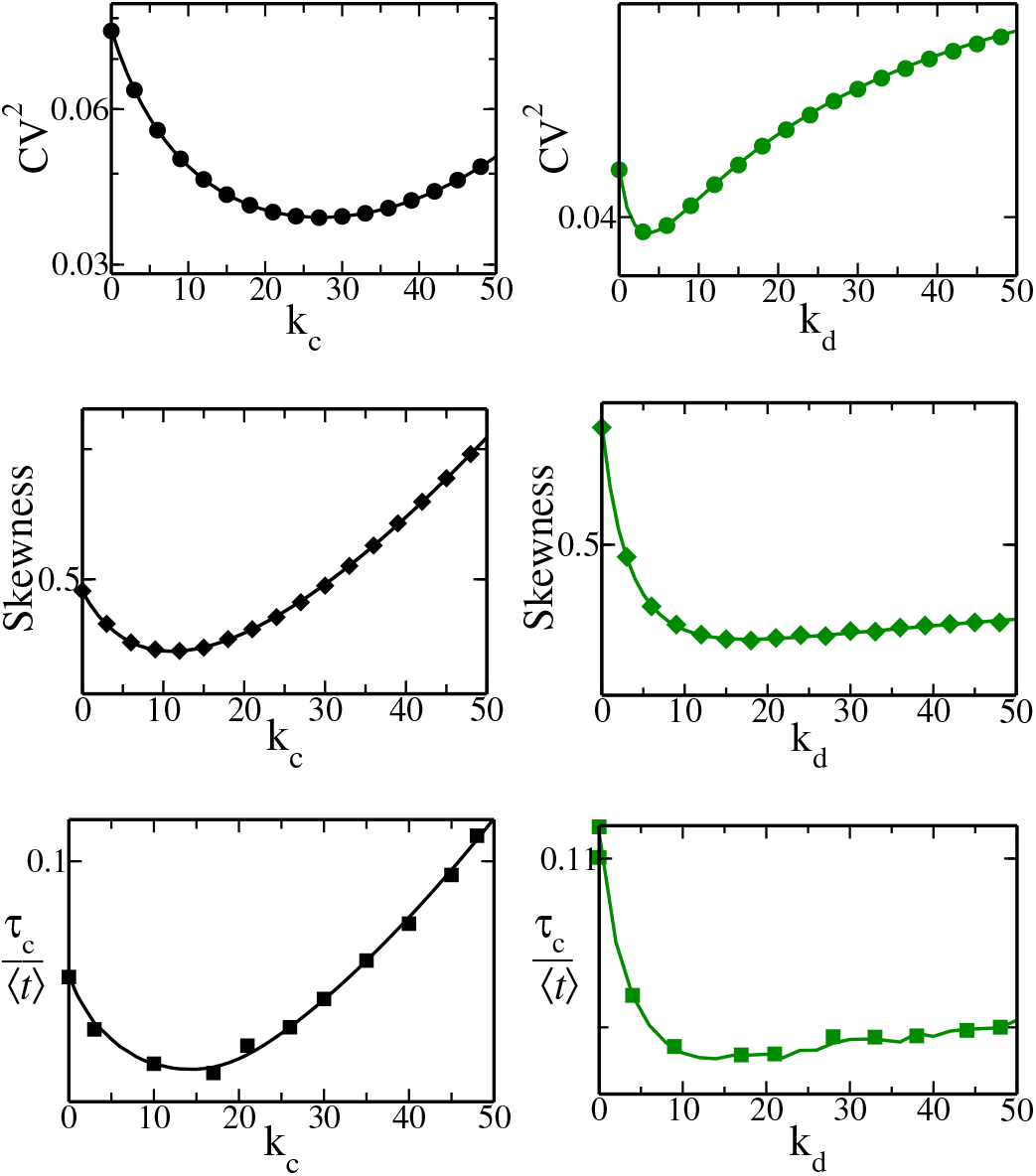
Model-II: Variation of *CV*^2^, Skewness and *τ*_*c*_*/*⟨*t* ⟩ with *k*_*c*_ (left panel, *k*_*d*_ = 1) and *k*_*d*_ (right panel, *k*_*c*_ = 37.5). Solid lines represent the analytically exact solutions, while the points with symbols correspond to KMC simulation data. The parameter set used for the KMC of the full Model-II is: *k*_*m*_ = 0.3, *k*_*p*_ = 24, *γ*_*m*_ = *γ*_*c*_ = 6, *X* = 100 and *γ*_*p*_ = 0.001. All rate parameters are in units of min^−1^.

From Eq. 13, we see that *b*^*e f f*^ monotonically increases and saturates with *k*_*d*_. Hence again due to the left panel of Fig 3, we are led to the exact prediction that the relative fluctuations of FPT in Model-II, will show non-monotonic dependence on *k*_*d*_ (as that changes *b*^*e f f*^). This is confirmed in the subfigures of the right panel of Fig. 4.

### D. Model-III: Variation of relative fluctuations with the switching parameters *k*_*c*_ and *k*_*d*_

The matter is a bit less straightforward in Model-III with positive post-transcriptional regulation. As we see in Eq. 14, now *b*^*e f f*^ as well as *k*_*m*_^*e f f*^ both increases monotonically with *k*_*c*_. Yet from Fig. 3, we see that the metrics of fluctuation behave qualitatively differently for *b*^*e f f*^ and *k*_*m*_^*e f f*^ – for the former, they are non-monotonic while for the latter they are monotonic. Hence it is hard to predict what would happen in Model-III. That is exactly what we find in the left panel of Fig. 5. While *CV*^2^ shows a shallow minimum, Skewness and *τ*_*c*_*/* ⟨*t*⟩ seems to be monotonic. Note that here too, theoretical *τ*_*c*_ is calculated using Eq. 11, while *τ*_*c*_ is calculated from the KMC data using Eq. 12.

**FIG. 5:**
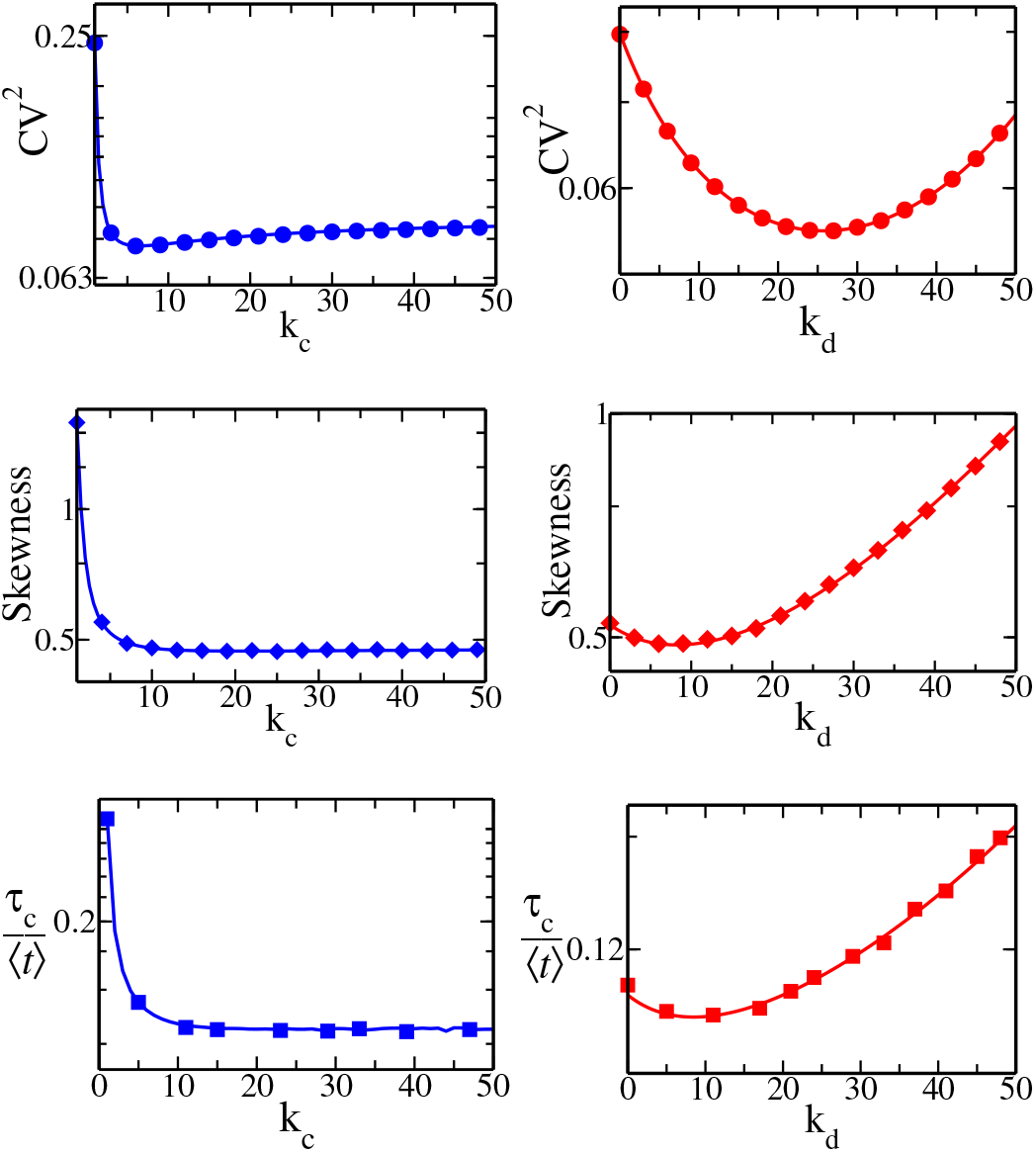
Model-III: Variation of *CV*^2^, Skewness and *τ*_*c*_*/*⟨*t*⟩with *k*_*c*_ (left panel, *k*_*d*_ = 5) and *k*_*d*_ (right panel, *k*_*c*_ = 7). Solid lines represent the analytically exact solutions, while the points with symbols correspond to KMC simulation data. The parameter set used for the KMC of the full Model-III is: *k*_*m*_ = 0.3, *k*_*pc*_ = 24, *γ*_*m*_ = *γ*_*c*_ = 6, *X* = 100 and *γ*_*p*_ = 0.001. All rate parameters are in units of min^−1^.

On the other hand, there is no dependence of *k*_*m*_^*e f f*^ on *k*_*d*_, while *b*^*e f f*^ monotonically decreases with *k*_*d*_. Hence it may be clearly concluded that the metrics of fluctuation will all have non-monotonic behaviour with *k*_*d*_ in Model-III, just based on their behaviour in Model-I (Fig. 3). We show that this expectation bears out and there is perfect match of theory and KMC data in Fig. 5 (right panel).

### E. Does post-transcriptional regulation increase or decrease relative fluctuations?

It is interesting to observe from Eq 13 that always *b*^*e f f*^ *< b* (because of the fractional factor multiplying *b*). Note that *b* = *k*_*p*_*/γ*_*m*_, so if we vary *b* by varying *k*_*p*_ without varying any other parameters in Model-II, there would be a lag of *b*^*e f f*^ from the value of changing *b* by a constant factor. As a result the *CV*^2^ for the regulated Model-II will be above the *CV*^2^ of the unregulated Model-I at small values of *b* (due to the lag of *b*^*e f f*^). This would change as *CV*^2^ of Model-I crosses its minimum. Now for larger *b*, the fluctuation of the regulated Model-II will go below that of unregulated Model-I (again due to the lag of *b*^*e f f*^). This crossing of the curves of relative fluctuation of the regulated versus unregulated model is an interesting result from our analytical arguments, which we confirm through simulations in Fig 6(a).

**FIG. 6:**
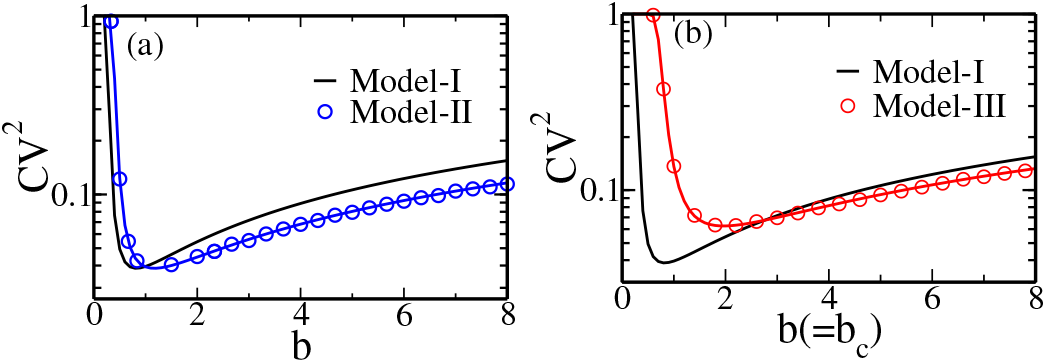
(a) The *CV*^2^ of Model-II goes from being above that of Model-I (at small *b*) to below it (at large *b*). The parameters in KMC of Model-II are *k*_*m*_ = 0.3, *k*_*c*_ = 3, *k*_*d*_ = 1, *γ*_*m*_ = *γ*_*c*_ = 6, *γ*_*p*_ = 0.001. (b) Similar crossing of *CV*^2^ curves of Model-III and Model-I as *b* is varied. We assume *b*_*c*_ of Model-III same as *b* of Model-I. The parameters in KMC of Model-III are *k*_*m*_ = 0.3, *k*_*c*_ = 5, *k*_*d*_ = 3, *γ*_*m*_ = *γ*_*c*_ = 6, *γ*_*p*_ = 0.001. All rate parameters are in units of min^−1^.

For Model-III, the relevant parameter is *b*_*c*_ and it can be compared to Model-I only if we assume the values *b*_*c*_ = *b*. If we do that from Eq 14, we see that *b*^*e f f*^ is again a fraction of *b*_*c*_. Thus again one expects a crossing of the curves of *CV*^2^, and this is exactly what we see in Fig 6(b).

### F. Is the U-shape in *CV*^2^ generic as a funtion of *k*_*c &*_ *k*_*d*_ ?

We showed above that the *CV*^2^ of FPTs as a function of binding (unbinding) rates *k*_*c*_ (*k*_*d*_) in sRNA regulated models, under the assumption of equal degradation rates of mRNA and mRNA-sRNA complex (i.e. *γ*_*m*_ = *γ*_*c*_), is U-shaped. In such cases, exact mappings and thus exact solutions were possible, which we could compare with KMC simulations. But in experiments this special condition may not generally apply. So it is important to relax this constraint and check using KMC simulations if this behaviour holds, even if we may not be able to provide an exact result to compare with. This is done in Fig. 7 by setting *γ*_*m*_ ≠ *γ*_*c*_. In the upper panel of the figure for Model-II, for two unequal sets of *γ*_*m*_ *& γ*_*c*_ (in filled and empty symbols) we see that *CV*^2^ has similar U-shapes as in Fig 4 with both *k*_*c*_ and *k*_*d*_. In the lower panel for Model-III we see that the minima are inconclusive with *k*_*c*_ but the U-shape is clearly visible with *k*_*d*_, again just as in Fig 5. Thus the exact results we found actually serve as very good guide to show a generic qualitative feature, namely the U-shape in *CV*^2^, which may be expected over a range of parameters *γ*_*m*_ *& γ*_*c*_ in experiments.

**FIG. 7:**
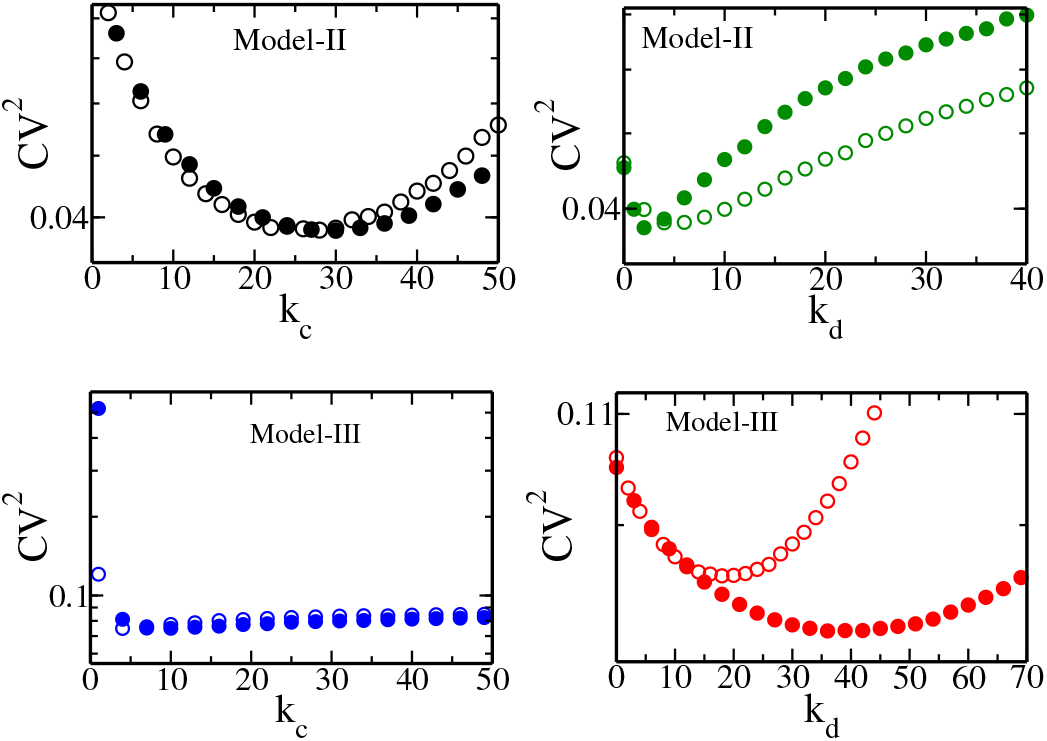
*CV*^2^ from KMC simulations as a function of *k*_*c*_ and *k*_*d*_ for *γ*_*m*_ =≠ *γ*_*c*_. Upper panel: for Model-II, *γ*_*c*_ = 4 (filled symbols) and *γ*_*c*_ = 8 (empty symbols) with *γ*_*m*_ = 6, *k*_*m*_ = 0.3, *k*_*p*_ = 24, *k*_*d*_ = 1, *γ*_*p*_ = 0.001. Lower panel: for Model-III, *γ*_*m*_ = 4 (filled symbols) and *γ*_*m*_ = 8 (empty symbols) with *γ*_*c*_ = 6, *k*_*m*_ = 0.3, 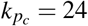, *k*_*c*_ = 37.5, *γ*_*p*_ = 0.001. All rate parameters are in units of min^−1^.

## IV. CONCLUSION

In the previous literature on theoretical analysis of sRNA regulation of gene expression, a useful mapping to the un-regulated case was found under assumptions of bursty protein production, and assuming the equality of degradation rates of mRNA and the mRNA-sRNA complex^68,69,76^. However, only linearized approximations were used to calculate the first passage statistics of protein threshold crossing, and compared with KMC simulations^68,69^. Now that exact solutions for the statistics of FPTs, for the unregulated problem are known^5,6,27^, we can exactly calculate the full distribution and other stochastic properties of protein threshold crossing times for proteins regulated by sRNAs. The exact full probability distribution, its moments and exponential tail have been explicitly obtained and presented here. Our KMC simulations show a perfect match with theory, in any region of parameter space which one may choose to study.

Moreover, due to the mappings, the sRNA regulated Models-II & III have the same theoretical expressions for metrics of FPT fluctuations as the unregulated model, only now with *effective* burst sizes *b*^*e f f*^ and transcription rates *k*_*m*_^*e f f*^. As a result, the interesting dependences on model parameters of the metrics of relative FPT fluctuations in models with sRNA regulation can now be studied exactly, without the assumptions required in the previous literature^68,69^. We particularly studied the dependence on the parameters *k*_*c*_ and *k*_*d*_, the association and dissociation rates of the mRNA-sRNA complex which are crucial measures of the strength of sRNA regulation. We would also like to stress that in the literature, *CV*^2^ is typically studied to understand fluctuations. Here we go beyond and also study Skewness (proportional to third order cumulant) and most importantly the characteristic time of the exponential tail of the distribution, which are measures of large deviation of the FPT distribution.

One interesting finding is that for negative regulation (Model-II), as a function of both *k*_*c*_ and *k*_*d*_ relative fluctuations of FPT are non-monotonic. Thus there are optimal values of *k*_*c*_ and *k*_*d*_ where relative fluctuations are a minimum. In molecular processes where the cost of fluctuations is high (such as stress responses) we can hypothesize that natural selection would have optimized the first passage time distribution to have minimum fluctuations. It would be very interesting to test this out in experiments. For example, if mutants with different mRNA, sRNA binding/unbinding abilities are experimentally created, the prediction of optimality that we make can be tested. For positive regulation (Model-III), the *CV*^2^ is non-monotonic as a function of *k*_*d*_, but as a function of *k*_*c*_ there is no clear U-shape. We have also shown that the U-shape of *CV*^2^ holds even when the constraint of equal degradation rates of mRNA and mRNA-sRNA complex is relaxed. It would be very interesting to perform experiments to test if this U-shaped *CV*^2^ can be observed in cells and if, in some pathways, sRNA regulation is optimized to minimize noise.

We also find that the relative fluctuations of the regulated Model-II in comparison to the unregulated one, go from high to low depending upon the size *b* of the protein burst. In case of Model-III, we see a similar trend if we set *b*_*c*_ = *b*.

The exact results that we present here reveal many interesting facts about the distribution of protein threshold crossing times. These results have been confirmed by carrying out KMC simulations, which show an exact match with the theoretical results. Our results can be used to motivate new experiments and data analysis procedures on the control of timing of molecular processes using sRNA regulation, as discussed above.

## ACKNOWLEDGMENTS

SYA acknowledges IIT Bombay for financial support through the institute post-doctoral fellowship.

## Appendix A: Model-II

If *m* denotes the number of mRNAs, *m*_*c*_, the number of mRNA-sRNA complexes, and *n* is the number of proteins, then the probability of finding *m, m*_*c*_ and *n* at time *t* is *P*(*m, m*_*c*_, *n, t*) and it follows the forward Master equation:

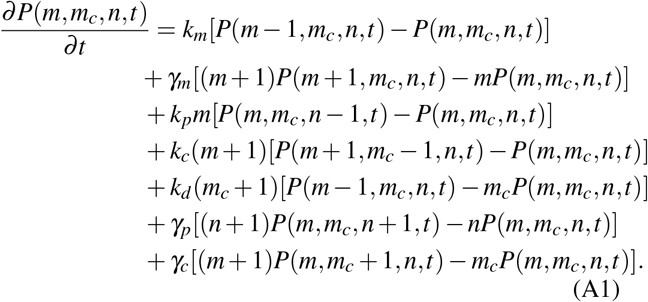

For the reader’s convenience, we reproduce some steps from^69^ to show how Model-II maps to Model-I in the *bursty* limit. Introducing the dimensionless parameters 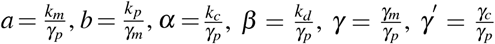 and by defining the generating function, 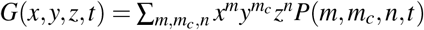, with *x* = *u* + 1, *y* = *v* + 1 and *z* = *w* + 1, Eq. A1 leads to:

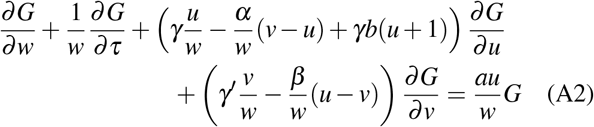

Eq. A2 can be solved by using the method of Lagrange characteristics^78^. Let *r* measure the distance along a characteristic. Then we have

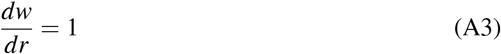

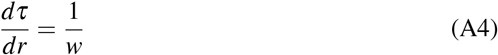

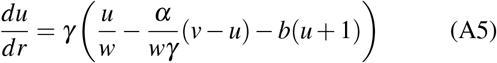

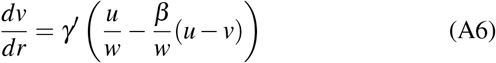

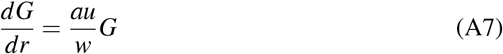

No, if we assume that

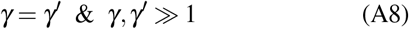

Then Eq. A5 and A6 solves for

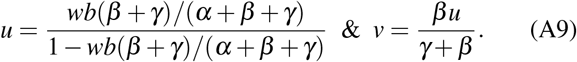

Substituting the *u* in Eq. A7 and using Eq. A3 one gets

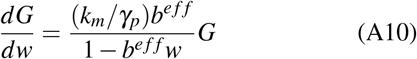

with 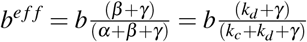.

## Appendix B: Model-III

If *m* is the number of mRNAs, *m*_*c*_ is the number of mRNA-sRNA complexes, and *n* is the number of proteins, then the Master equation of Model-III is^67^

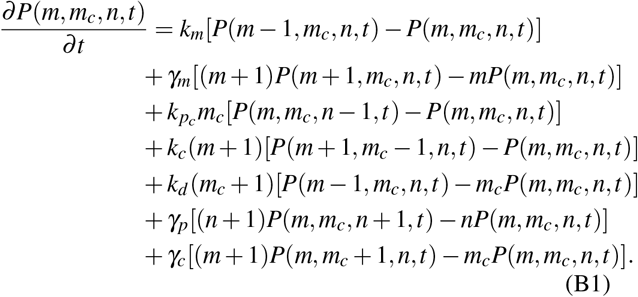

We reproduce some steps from^67^ to show how Model-III maps to Model-I in the *bursty* limit. Introducing the dimensionless parameters and the generating function *G* as in Model-II and 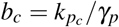, one can convert Eq. B1 into a first-order partial differential equation:

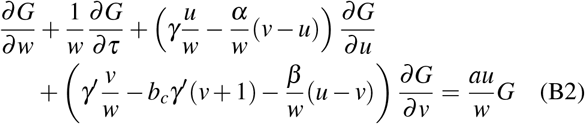

Eq. B2 can be solved by using the method of Lagrange characteristics^78^. If *r* measures the distance along a characteristic, then we get

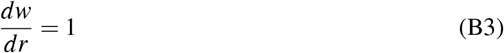

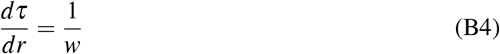

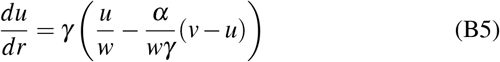

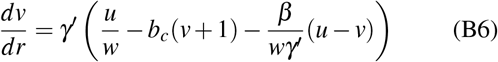

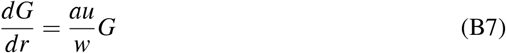

Now if assumptions as in Eq. A8 are true then Eq. B5 and B6 solves for

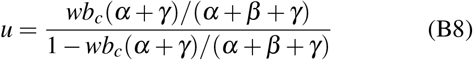

Substituting the *u* in Eq. B7 and using Eq. B3 one gets

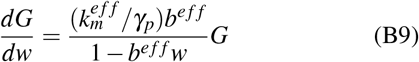

with 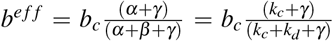 and 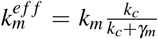 Where 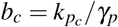.

## Appendix C: Model-IV

In this model, protein translation occurs from both mRNA and the mRNA-sRNA complex. Let *m* be the number of mRNAs, *m*_*c*_ the number of mRNA-sRNA complexes and *n* the number of proteins, then the Master equation of Model-IV is^69^

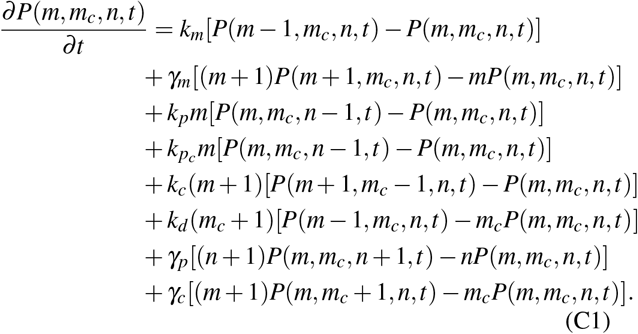

We reproduce some steps from^69^ to show how Model-IV maps to Model-I in the *bursty* limit. Introducing the dimensionless parameters and the generating function *G* as in Model-II and Model-III, one can convert Eq. C1 into

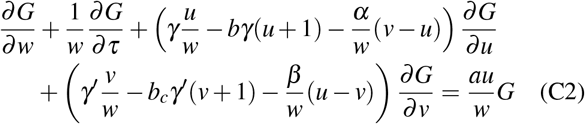

Eq. C2 can be solved by using the method of Lagrange characteristics^78^. If *r* measures the distance along a characteristic, then we get

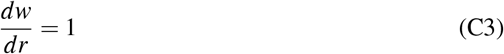

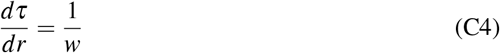

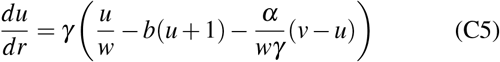

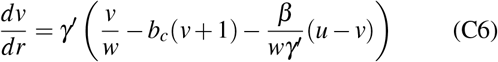

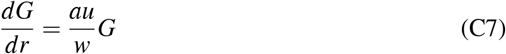

Now, if we assume that

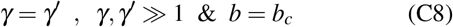

then Eq. C5 and C6 solves for

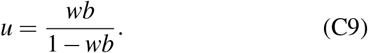

Substituting the *u* in Eq. C5 and using Eq. C7 one gets

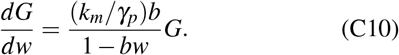

Here, *b*^*e f f*^ = *b* and 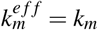.

